# FERARI: a novel tether involved in endocytic recycling

**DOI:** 10.1101/500082

**Authors:** Jachen A. Solinger, Harun-Or Rashid, Anne Spang

## Abstract

Endosomal transport is essential for cellular organization and compartmentalization and cell-cell communication. Sorting endosomes provide a crossroad for various trafficking pathways and determine recycling, secretion or degradation of proteins. The organisation of these processes requires membrane tethering factors to coordinate Rab GTPase function with membrane fusion. Here, we report a conserved tethering complex that acts in the Rab11 recycling pathways at sorting endosomes, which we name FERARI (Factor for Endosome Recycling And Retromer Interactions). The Rab binding module of FERARI consists of Rab11FIP5/RFIP-2 and rabenosyn-5/RABS-5, while the SNARE interacting module comprises VPS45 and VIPAS39/SPE-39. Unexpectedly, the membrane fission protein EHD1/RME-1 is also a FERARI component. Thus, FERARI appears to combine fusion activity through the SM protein VPS45 with pinching activity through EHD1/RME-1 on SNX-1-positive endosomal membranes. We propose that coordination of fusion and pinching through a kiss-and-stay mechanism drives sorting at endosomes into recycling pathways.

## Introduction

Endocytosis is an essential process in the communication of the cell with its environment, controlling cell growth and regulating nutritional uptake. About 70-80% of endocytosed material is recycled back from sorting endosomes to the plasma membrane through a variety of pathways. Defects in recycling lead to a myriad of human diseases such as cancer, ARC (Arthrogryposis–renal dysfunction–cholestasis), BBS (Bardet–Biedl syndrome) or Alzheimer’s disease ^1–8^. In spite of their importance, sorting into recycling endosomes and the interaction of recycling with sorting endosomes remain still poorly understood.

The identity of recycling endosomes is largely characterized by the presence of a specific Rab GTPase on the limiting membrane of the organelle. For example, Rab4 and Rab11 are present on different types of recycling endosomes directly targeted to the plasma membrane, while Rab9 is involved in recycling from the sorting endosome to the TGN and Rab10 has multiple reported roles in recycling from endosomes to the TGN or basal lateral membranes and exocytosis from the TGN ^9, 10^. The different recycling pathways also highlight a robustness of the system, in that if one recycling pathway is impaired, alternative routes are still available. Yet, it also underscores the difficulty in studying these pathways because of their redundancy.

The number of recycling pathways is reduced in simpler eukaryotes such as yeast and *C. elegans*, making them prime systems to elucidate the basic mechanisms. *C. elegans* lacks clear homologues of Rab9 and Rab4. Yet, it is able to maintain complex organ structures like polarized intestinal epithelia cells, which provide an excellent system to analyze recycling processes ^11, 12^. In fact, the first recycling protein, RME-1, was discovered in *C. elegans* ^13^. RME-1 and the homologous human EHD proteins were shown to have tubulation and vesiculation activities ^14–16^. EHD1 interacts with Rab11 effectors Rab11FIP2 and Rab11FIP5 and is involved in Rab11-dependent recycling from tubular endosomal networks ^17, 18^. The formation of such tubular structures requires the action of BAR domain proteins, most prominently sorting nexins (SNXs). Indeed, SNX4 associates with Rab11 recycling endosomes ^19, 20^. In addition, EHD1 interacts with rabenosyn-5 to form recycling compartments ^21^.

Besides the coordination of recycling pathways, sorting endosomes have to cope with incoming material from the TGN through the biosynthetic route. This cargo has to be sorted into different domains presumably overlapping with the recycling pathways. Fusion of transport containers with endosomes is regulated by tethering factors and SNAREs. CORVET and HOPS tethering factors have been demonstrated to act on early endosomes and late endosomes, respectively ^12, 22–25^. The yeast HOPS and CORVET complexes only differ in two subunits, which recognize Rab7/Ypt7 and Rab5/Vps21, respectively ^22^. In metazoans, two of the shared HOPS/CORVET components are duplicated. Vps16 has a homolog, VIPAS39/SPE-39 and the Sec1/Munc18 (SM) protein Vps33 is present as Vps33A/VPS-33.1 and Vps33B/VPS-33.2 ^24^. Moreover, all eukaryotes contain another SM protein, VPS45, with a proposed function together with rabenosyn-5 in endosomal recycling ^26^.

We had speculated recently that VIPAS39/SPE-39 might be part of two HOPS/CORVET types of complexes, one in conjunction with Vps33B/VPS-33.2 and one with VPS45, which we named CHEVI (class <u>C H</u>omologs in <u>E</u>ndosome-<u>V</u>esicle Interaction) and FERARI (<u>F</u>actors for <u>E</u>ndosome <u>R</u>ecycling <u>A</u>nd <u>R</u>etromer Interaction), respectively ^27^. Here, we demonstrate the existence of FERARI in *C. elegans* and mammalian cells. FERARI plays a key role in Rab11-dependent recycling from sorting endosomes to the plasma membrane. We propose a model in which FERARI supports kiss-and-stay of Rab11-positive structures on tubular sorting endosomes to aid the formation of recycling endosomes.

## Results

### FERARI is a conserved complex

We have recently postulated the existence of a FERARI tethering complex containing VIPAS39/SPE-39 and the SM protein VPS-45, which we named FERARI ^27^. To test whether such a complex occurs, we co-expressed *C. elegans* GST-tagged SPE-39 and His-VPS-45 in *E. coli*. His-VPS-45 was co-purified with SPE-39-GST (Fig. 1 A). VPS-33.2, which together with SPE-39 is part of the CHEVI complex ^27, 28^ served as a positive control, while the HOPS specific VPS-33.1 was unable to bind to SPE-39. These data indicate that SPE-39 and VPS-45 can indeed directly interact. This interaction was confirmed by yeast-two-hybrid interaction and pull-down experiments. Moreover, FERARI is conserved in mammalian cells, as VPS45 was pulled-down by VIPAS-39 (Fig. 1C). VPS45 has been shown to directly interact with rabenosyn-5/RABS-5 ^29, 30^. Therefore, we asked whether SPE-39 could bind RABS-5. Similar to VPS45, rabenosyn-5/RABS-5 interacted with VIPAS/SPE-39 (Fig. 1B-E, Fig. S1A, E and G).

**Figure 1:**
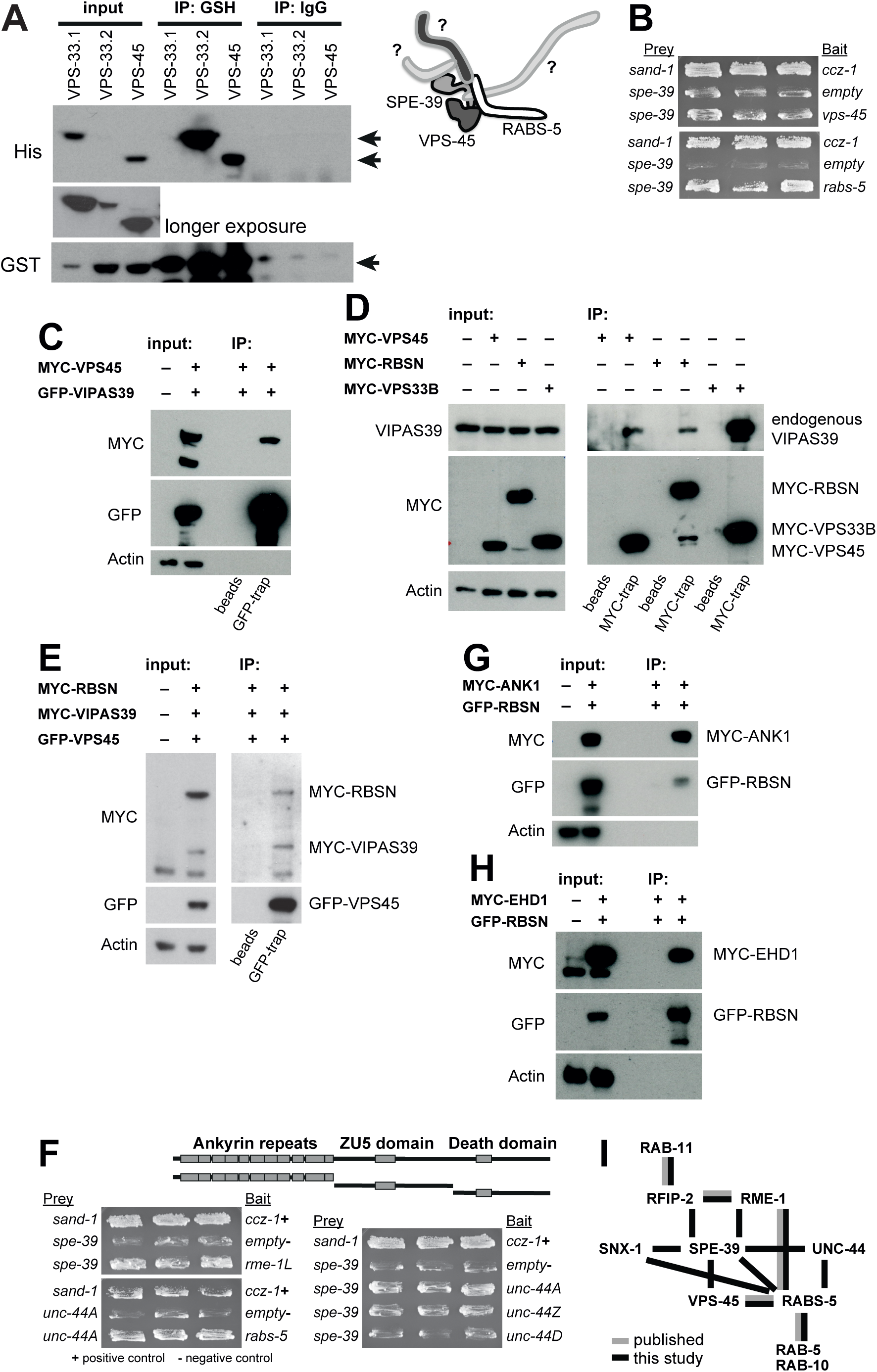
Conserved interactions between FERARI components. (A) Recombinant SPE-39 protein from *E. coli* interacts with VPS-45. SPE-39-GST was co-expressed with VPS-33.1, VPS-33.2 or VPS-45. Pull-downs were performed with either GSH-or IgG-beads as indicated. Full-length SPE-39-GST was expressed and could be bound to the beads. VPS-33.2 bound very stably to SPE-39 and was used as a positive control (see upper arrow on the right, VPS-33.1 and VPS-33.2 have a very similar size), while VPS-33.1 (a subunit of the HOPS complex) did not interact even while the expression was higher than for VPS-33.2. Interaction between SPE-39 and VPS-45 was also observed (see lower arrow on right). Schematic representation shows the initially known FERARI components that were used to bind additional interactors. (B) Yeast two hybrid assays show interactions between SPE-39 and VPS-45 as well as SPE-39 and RABS-5. Three independent yeast transformants were grown on plates with galactose but lacking leucine. The previously described interaction between SAND-1 and CCZ-1 was used as positive control, empty vector of activation domain plasmid pJG4-5 as negative control. (C) Interaction between VIPAS39 to VPS45 in HEK-293 cells. Cells were co-transfected with myc-tagged VPS45 and GFP-tagged VIPAS39 for 48 h. Lysates prepared from transfected cells were subjected to immunoprecipitation with either control beads (only agarose beads as negative control) or GFP-trap beads. Immunoprecipitated proteins and inputs were then probed with antibodies against myc, GFP and actin. n= 5 independent experiments. (D) Interaction of endogenous VIPAS39 with rabenosyn-5 and VPS45. HEK-293 cells were transiently transfected with myc tagged Rabenosyn-5, VPS45, or VPS33B and lysates were prepared. Immunoprecipitation was performed with myc-trap and control beads and immunoblotted with VIPAS39, myc and actin antibodies. Interaction between VIPAS39 and VPS33B served as a positive control. n= 3 independent experiments (E) Interaction of VIPAS39, VPS45 and rabenosyn-5 in triple transfected cells. Lysates were prepared from GFP-VIPAS39, myc-Rabenosyn-5 and myc-VPS45 triple transfected HEK-293 cells. GFP-VIPAS39 was immunoprecipitated from cell lysates with GFP-trap beads, followed by detection of co-immunoprecipitated myc-Rabenosyn-5 and myc-VPS45. n= 3 independent experiments (F) Yeast two hybrid interactions between SPE-39 and RME-1, UNC-44, as well as between UNC-44 and RABS-5. Yeast was transformed with the DBD-plasmid pEG202 and the AD-plasmid pJG4-5 bearing the indicated genes and grown on plates lacking leucine. UNC-44 was subcloned into three domains as indicated in the schematic drawing. Interaction was through the ankyrin repeats. (G and H) Binding of ANK1 and EHD1 to rabenosyn-5 (RBSN) in HEK-293 cells. GFP-Rabenosyn-5 was transiently co-expressed with myc-ANK1 (G) and myc-EHD1 (H), and GFP-Rabenosyn-5 was immunoprecipitated with GFP-trap. Immunoprecipitated proteins were detected using myc and GFP antibodies. Actin served as a loading control. n= 3 independent experiments (I) Summary of interactions between FERARI components. This includes all interactions seen in Y2H and pull-downs with worm and human proteins and previously published data.

Based on the multisubunit structures of the HOPS and CORVET complexes, we speculated that the FERARI complex might contain more subunits (Fig. 1A). To identify additional components, we incubated the bacterially expressed SPE-39/VPS-45 complex with worm lysate and performed mass spectrometry (Table S1). In a secondary screen, we performed yeast-two-hybrid assays and pull-downs from selected, streamlined candidates. This analysis revealed two new components of the FERARI complex: the Epsin-homology domain (EHD) containing protein RME-1 and UNC-44 (Fig. 1F).

UNC-44 contains ankyrin motifs and a death domain. While SPE-39 interacted well with RME-1 and the ankyrin motifs of UNC-44, it did not bind to the death domain (Fig. 1F, Fig. S1C and D). Both, RME-1 and UNC-44 are conserved in mammals. RME-1 is homologous to EHD1-4, and UNC-44 to ANK1-3. Since EHD1 has been shown to act at endosomes and EHD2 at the plasma membrane ^31, 32^, we focused on EHD1. EHD1 co-precipitated and co-localized with rabenosyn-5 (Fig. 1H and Fig. S2A and B). Much less is known about ANK1-3. ANK1 co-precipitated with rabenosyn-5, but showed rather limited co-localization with rabenosyn-5, indicating that it might be a more dynamic subunit of the complex or the other ANKs could also take part in the complex (Fig. 1G and Fig. S2A). Indeed, ANK3 appears to be the ANK protein most highly expressed in HeLa cells (Fig. S2D). Thus, a combination of biochemistry, co-immunoprecipitation and yeast-two-hybrid assays revealed the existence of a conserved FERARI tethering complex, consisting of VIPAS39/SPE-39, VPS45, rabenosyn-5/RABS-5, EHD1/RME-1 and ANK1/ANK3/UNC-44 with a proposed architecture illustrated in Fig. 1I, based on our own data (see also below) and on literature ^17, 18, 21, 29, 30^.

### Loss of FERARI affects endosomal recycling

VPS45, rabenosyn-5 and EHD1 have been implicated in recycling from sorting endosomes in mammalian cells ^30, 32^. To test whether FERARI might be involved in recycling, we knocked out VIPAS39, VPS45 and rabenosyn-5 in HeLa cells using CRISPR-Cas9 (Fig. S2C) and expressed GFP-Rab11 as a recycling marker in those cells (Fig. 2A). As ANK3 appeared to be the major ankyrin protein in HeLa cells (Fig. S2D), we knocked out ANK3 (Fig. S2C). In FERARI knockout cells, Rab11-positive structures were enlarged when compared to control cells (Fig. 2A and B). We went back to our animal model to confirm and explore further the observed phenotype, and to investigate whether FERARI is organ-specific or functions ubiquitously. The *C. elegans* intestine is a tubular epithelium with a pair of juxtaposed cells forming a tube, the apical gut lumen (Fig. 2C). This system provides an excellent system to study membrane traffic in the metazoa ^12,33–36^.

**Figure 2:**
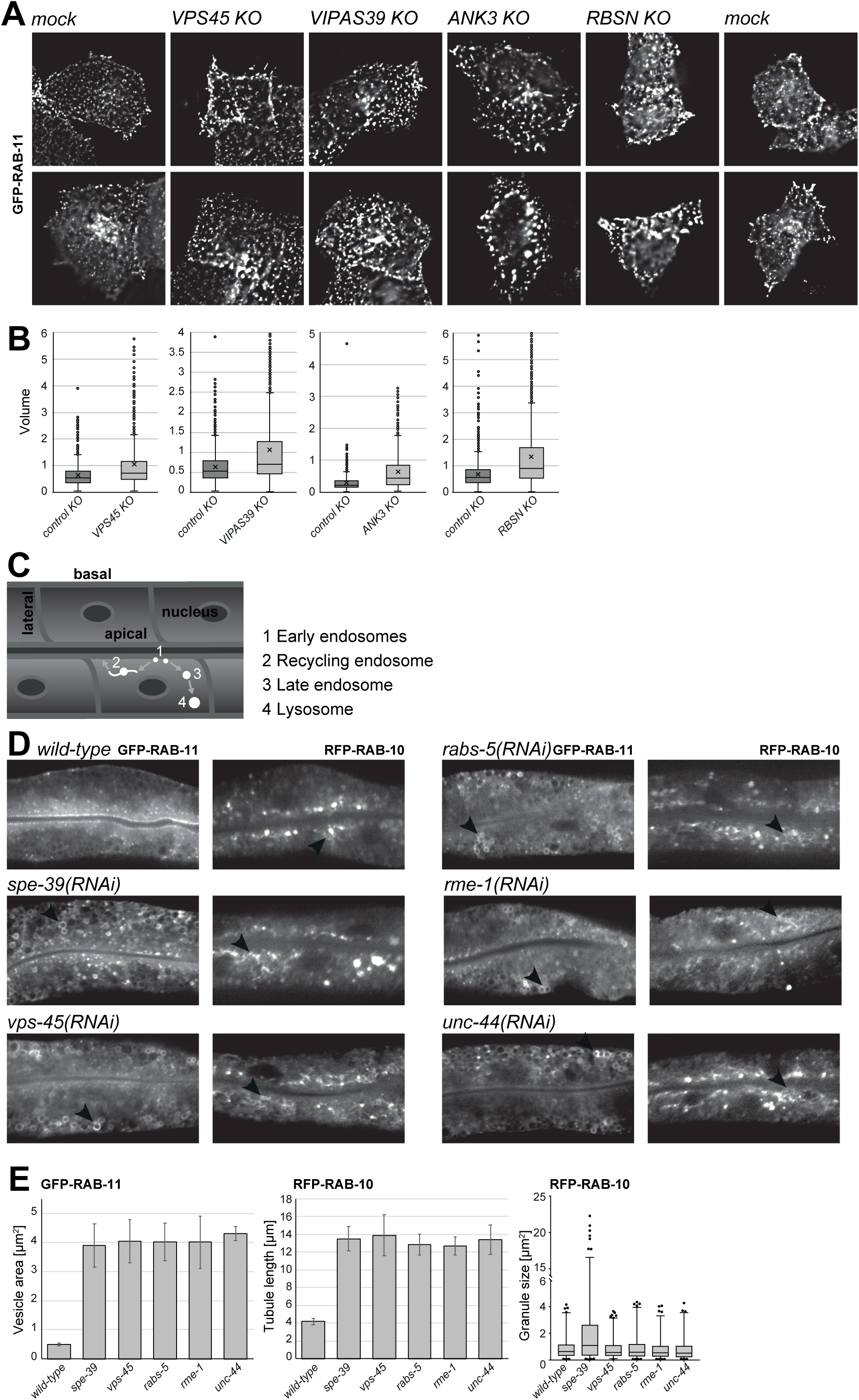
Loss of FERARI affects endosomal recycling. (A) CRISPR-Cas9 knock-outs of vipas39, vps45, rbsn and ank3 cause enlargement of Rab11 compartments in HeLa cells. Representative images from live cell imaging of Rab11-GFP in control and indicated knock-out cells. (B) Quantification of deconvolved and FIJI processed images shown in (A). n= 45-60 cells from two-three biologically independent experiments. Sizes are given in arbitrary units showing an enlargement of the compartments compared to mock treated cells. (C) Schematic representation of *C. elegans* intestinal cells and approximation of different compartments in the endosomal system (shown in white). (D) GFP-RAB-11 structures are changed by *FERARI(RNAi)* (D left side). Normal RAB-11 structures near the apical membrane (in *wild-type)* are greatly enlarged and mis-localized to more basal positions inside intestinal cells (indicated by arrowheads in D). RFP-RAB-10 compartments are also strongly affected by RNAi of FERARI components (D right side). The *wild-type* round compartments with small networks (see arrowhead) are changed to grossly enlarged networks in FERARI knock-down worms (arrowheads for *spe-39, vps-45, rabs-5, rme-1* and *unc-44* knock-downs). The enlarged globular compartments in *spe-39(RNAi)* worms (right panel) might be explained by an additional role of SPE-39 in the CHEVI complex. (E) Quantification of RAB-11 and RAB-10 phenotypes shown in (D). RAB-11 compartments and RAB-10 tubules were measured in 6 worms (10 structures each), RAB-10 globular compartments were quantified semi-automatically and comprise 50-120 structures (per worm) in 6 different worms.

We used *C. elegans* animals expressing GFP-RAB-11 and RFP-RAB-10, markers for apical and baso-lateral recycling compartments, respectively ^37, 38^. In mock treated animals, RAB-11 was mostly localized at the apical cortex, while RAB-10 was present on more internals structures (Fig. 2D). Similar to what we had observed in mammalian cells, loss of FERARI caused the enlargement of RAB-11-positive structures (Fig. 2D and E), indicating that FERARI function is conserved. However, not only apical recycling was perturbed as the tubular part of the RAB-10 compartment was largely extended, while the size of the granular RAB-10 structures remained unchanged (Fig. 2D and E). Human transferrin receptor (hTfR) has been used as a cargo to study recycling in the *C. elegans* intestine ^38^. Knockdown of FERARI led to a strong reduction of the hTfR signal in the gut, suggesting that it less efficiently sorted into recycling endosomes and therefore degraded in the lysosome (Fig. S3A). We conclude that FERARI plays a role in both apical and baso-lateral recycling.

*C. elegans* oocytes provide another excellent model for endocytosis. Yolk is taken up by the maturing oocytes via receptor-mediated endocytosis. Recycling of the yolk receptor RME-2 is essential for efficient yolk uptake. Knock-down of FERARI reduced the level of yolk in oocytes (Fig. S3B and C). This reduction was due to impaired recycling of RME-2, which accumulated in internal structures (Fig. S3D). Thus, our data indicate that FERARI is involved in recycling to the plasma membrane in different tissues.

### Rab11FIP5/RFIP-2 is part of FERARI and provides the link to Rab11

We observed a strong defect upon loss of FERARI function on Rab11 recycling compartments. None of the FERARI components has been shown to directly bind Rab11. However, EHD1 was shown to bind the Rab11 effectors Rab11FIP2 and Rab11FIP5 ^18^ in mammalian cells. To assess, whether the Rab11FIPs could interact with the thus far identified FERARI components, we performed a GFP-pull down in HEK 293 cells expressing GFP-rabenosyn-5 and MYC-Rab11FIP2 or MYC-Rab11FIP5 (Fig. 3A). While Rab11FIP5 was co-precipitated, Rab11FIP2 was not, indicating that Rab11FIP5 is the link between Rab11 and FERARI. We also confirmed that Rab11FIP5 indeed bound to Rab11 (Fig. 3B). To corroborate that Rab11FIP5 was the RAB interaction module, we co-expressed VIPAS39 and RAB11 with or without Rab11FIP5 (Fig. 3C). While VIPAS39 was barely detected in a RAB11-GFP pulldown, this interaction was substantially stronger in the presence of Rab11FIP5, suggesting that Rab11FIP5 could bridge the interaction between VIPAS39 and RAB11. When we knocked out Rab11FIP5 in HeLa cells, Rab11-positive structures were enlarged, similarly to what we observed for the other FERARI components (Fig. 3D and E; compare to Fig. 2A). Consistently, Rab11FIP5 co-localized with rabenosyn-5 (Fig. S2A and B). In *C. elegans*, there is only one homolog of Rab11FIP2 and Rab11FIP5, the so far uncharacterized Y39F10B.1, which we named RFIP-2. Knockdown of RFIP-2 phenocopied the effect of the FERARI(RNAi) in the *C. elegans* intestine (Fig. 3F and G). Moreover, RFIP-2 interacted with FERARI components in pull-downs (Fig. 3H). Therefore, we propose that Rab11FIP5/RFIP-2 represents one branch of the RAB interaction module of FERARI (Fig. 1I).

**Figure 3:**
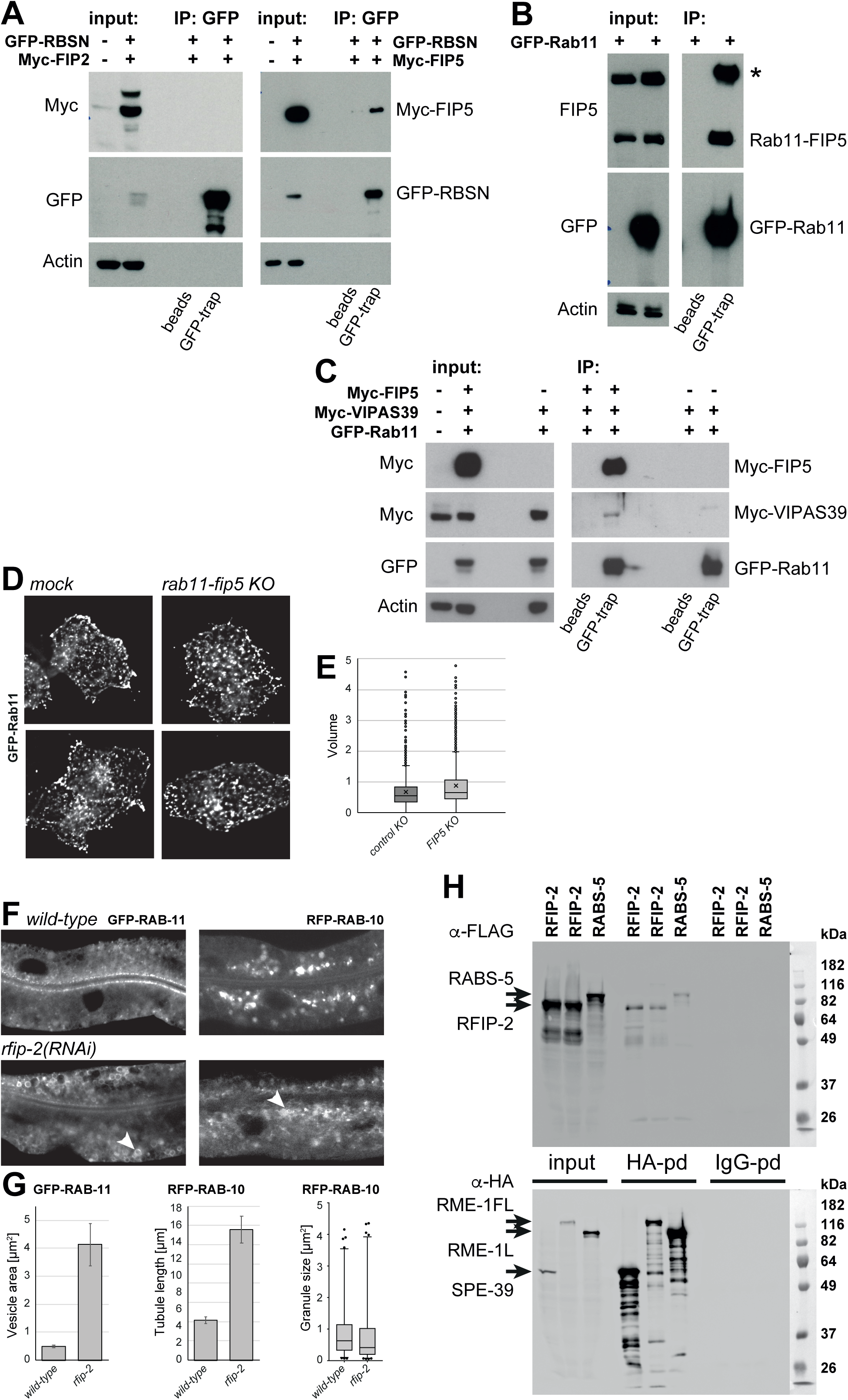
Rab11FIP5/RFIP-2 is part of FERARI and provides the link to Rab11. (A) Rabenosyn-5 binds to Rab11FIP5 but not to Rab11FIP2 in HEK-293 cells. myc-tagged Rab11FIP5 but not Rab11FIP2 was co-immunoprecipitated with GFP-Rabenosyn-5. Representative images from 3 independent experiments. (B) Western blot of immunoprecipitation of endogenous Rab11FIP5 with GFP-Rab11 in HEK-293 cells. (C) Rab11 binds to FERARI through Rab11FIP5. HEK-293 cells were co-transfected with GFP-Rab11 and myc-VIPAS39 in the presence or absence of myc-Rab11FIP5. Interaction between VIPAS39 and Rab11 was enhanced in the presence of Rab11FIP5. (D) Live cell imaging of Rab11FIP5 CRISPR/Cas9 knock-out HeLa cells. Rab11 compartments were enlarged, similar to the phenotype of the knockout other FERARI subunits. (E) Quantification of deconvulated and FIJI processed images shown in (D). About 2,300 GFP-Rab11 positive particles were analyzed from 50 cells from two independent experiments. (F) Phenotypes of *rfip-2* knock-downs in worm intestinal cells with GFP-RAB-11 (left) and RFP-RAB-10 (right) markers. Phenotypes are the same as for other FERARI components (see Fig. 2D), namely enlarged globular RAB-11 compartments and enhanced networks of RAB-10 (indicated by arrowheads). (G) Quantification of *rfip-2* phenotypes shown in (F). (H) Interactions between proteins in the yeast two hybrid vectors were tested biochemically with pull-downs, because no growth was seen on plates lacking leucine. Worm RFIP-2 (Y39F10B.1) binds to RME-1 and SPE-39. Interaction between RME-1 and RABS-5 also occurs in worms.

### FERARI may act on sorting endosomes but is not essential for transport to the lysosome

Our data so far suggest that FERARI possibly functions on sorting endosomes. If this interpretation is correct, we might be able to detect an effect on the localization of the Rab proteins found on sorting endosomes, RAB-5 and RAB-7. Upon silencing of FERARI components, the RAB-5 signal appeared to be more concentrated at the apical membrane and also part of the RAB-7 signal became enriched in the same area in FERARI(RNAi) compared to mock-treated animals (Fig. 4A and B). One possible explanation for this finding is that reducing recycling capacity could prematurely activate Rab conversion from RAB-5 to RAB-7 positive membranes. If this assumption was correct, disruption of Rab conversion should restore the RAB-5 localization. SAND-1 is part of a RAB-7 GEF complex and a key regulator of Rab conversion ^39–41^. Concomitant loss of SAND-1 and FERARI restored RAB-5 localization (Fig. 4C and D). Surprisingly, RAB-7 positive compartments were also partially restored under these conditions, even also moderately rescuing the transport defect of hTfR in *sand-1 -/-* animals (Fig. 4C-E). Likewise, the *sand-1 -/*-transport block in coelomocytes, which are scavenger cells of the *C. elegans* innate immune system, was also partially rescued (Fig. S4). Thus, loss of FERARI impacts RAB-5 and RAB-7-positive endosomes, consistent with a function of FERARI on sorting endosomes.

**Figure 4:**
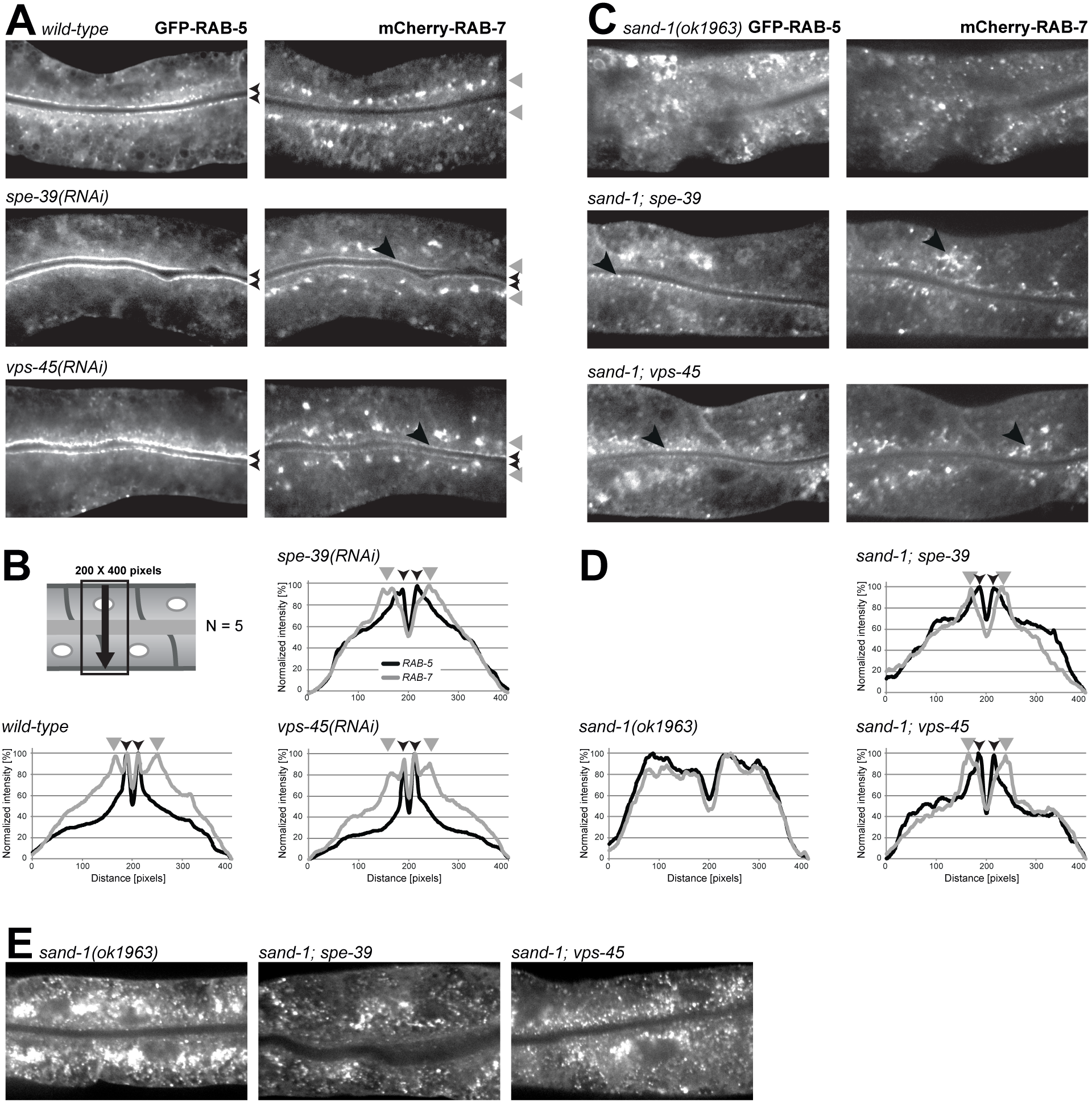
FERARI may act on sorting endosomes but is not essential for transport to the lysosome. (A) RAB-5 positive compartments are affected by *spe-39(RNAi)* and *vps-45(RNAi)*. GFP-RAB-5 signal remains confined to the apical membrane and these compartments even seem to mature to RAB-7 (at least partially, as seen by the slight mCherry-RAB-7 signal at the apical membrane, arrows in right panels). General RAB-7 compartment is largely unaffected (see gray arrowheads). (B) Quantification of RAB-5 and RAB-7 phenotypes. Schematic representation of a worm gut and the area used for quantification (line plot with 200 pixels width). Small arrows indicate RAB-5 peaks (also the lines along the apical membrane in the worm intestines in A), arrowheads in gray indicate normal RAB-7 compartments (also position in the worm intestine in A). (C) Enlarged RAB-5 compartments in *sand-1(ok1963)* worms are partially restored in *spe-39* and *vps-45* knock-downs (arrowheads in right panels). The dispersed and disorganized RAB-7 compartments of *sand-1(ok1963)* worms are also partially suppressed by loss of FERARI (right panels). (D) Quantification of *sand-1* suppression phenotypes as in (B). Since compartments in *sand-1(ok1963)* worms are mis-localized, the average distribution in 5 worms is a broad peak across the whole intestinal cell. In *spe-39(RNAi)* and *vps-45(RNAi)* the peaks of RAB-5 and RAB-7 look more similar to *wild-type* (B). (E) In *sand-1(ok1963)* worms the degradation pathway is blocked and hTfR-GFP accumulates in very bright endosomes. Knock-down of *spe-39* and *vps-45* cause less hTfR-GFP signal, because in the absence of recycling the degradative pathway is the only alternative available for hTfR-GFP which will lead to more degradation in these worms.

### FERARI is present on and required for the maintenance of SNX-1 active recycling compartments

The sorting nexins SNX1 and SNX4 in mammals and the *C. elegans* homolog SNX-1 are present on the tubular part of sorting endosomes ^42, 43^. Mild overexpression of mCherry-SNX-1 slightly increases the size of tubular portions of sorting endosomes (Fig. 5A), while stronger overexpression of GFP-SNX-1 caused extensive tubulation in intestinal cells and could not be used for these analyses (data not shown). To test if the SNX-1 positive tubules are active recycling compartments, we blocked apical recycling by *rab-11(RNAi)* (Fig. 5A). As expected, the SNX-1 tubules were extended to long networks when RAB-11 was depleted. This finding suggests that RAB-11 drives recycling through these tubules and that the SNX-1 tubules are dynamic structures that respond to recycling flux. Moreover, RAB-5, RAB-7 and RAB-11 co-localized with SNX-1 tubules to a varying degree (Fig. 5B and C, movies S1-S4), consistent with the notion that SNX-1 positive structures represent active recycling compartments on sorting endosomes. Of note, RAB-7 occupied larger domains on the SNX-1 tubules, while the RAB-11 signal was more vesicular and tubular with a smaller contact site. The Golgi marker MANS-1 and the lipid droplet marker DHS-3 did not show any appreciable co-localization with SNX-1 tubules (Fig. 5B and C, movies S6 and S7). Given the ease of detecting the SNX-1 tubular structure, in the *C. elegans* intestine, we focused on *C. elegans* to elucidate further the function of FERARI.

**Figure 5:**
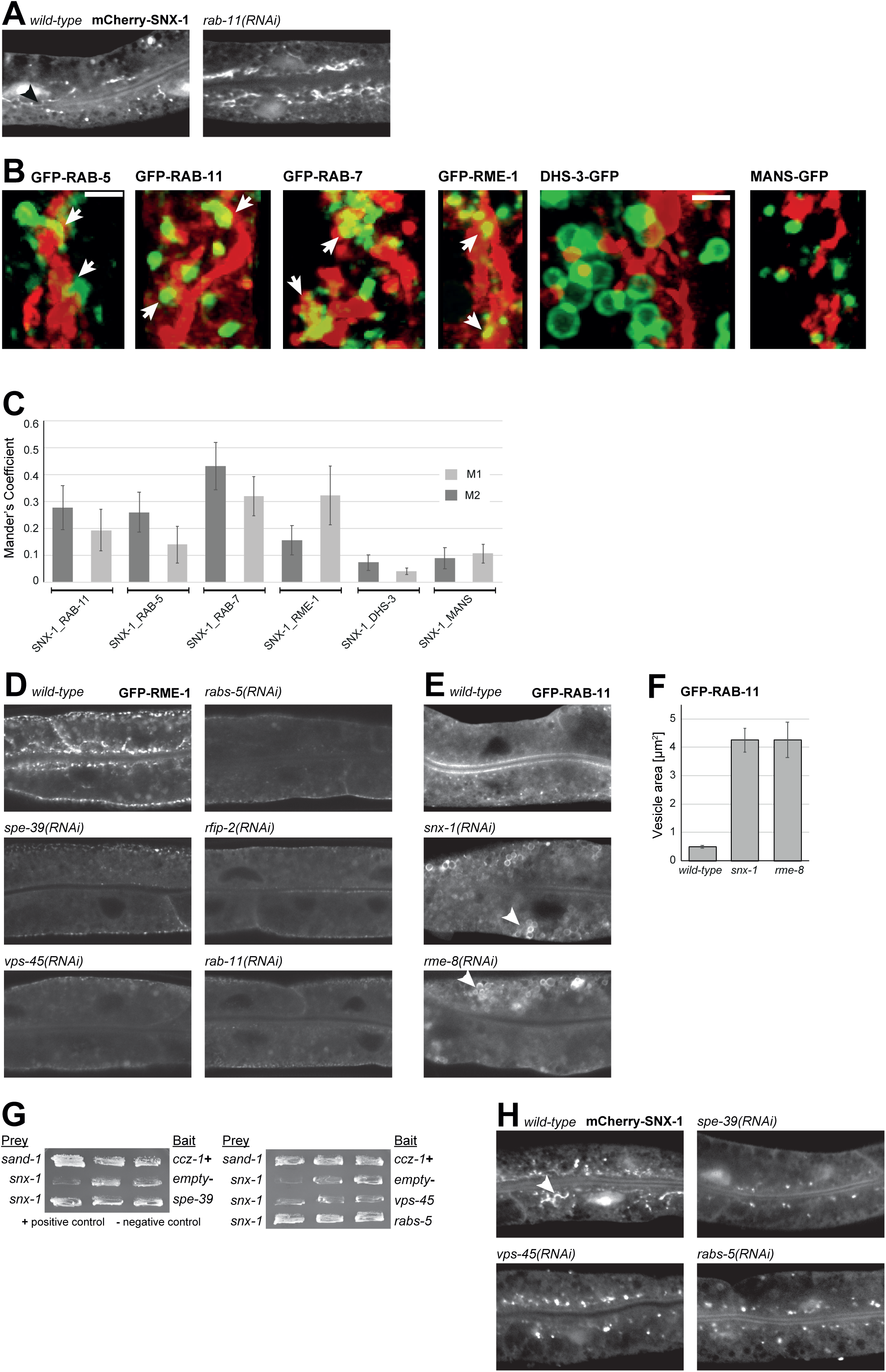
FERARI is present on and required for the maintenance of SNX-1 active recycling compartments. (A) Sorting networks can be observed using an mCherry-SNX-1 marker (arrow). SNX-1 tubules are elongated and accumulate in *rab-11(RNAi)* worms. (B) mCherry-SNX-1 networks are relatively short and sometimes coincide with the position of RAB-11, RAB-5, RAB-7 and RME-1 compartments. Worms with double markers mCherry-SNX-1 and either RAB-5, RAB-11, RAB-7 and RME-1 show that some overlap exists between the networks and the globular compartments (arrows in B). As controls with very little to no overlap DHS-3 (lipid droplets) and MANS (Golgi) were used. (C) Quantification of co-localization using Mander’s coefficient. (D) Localization of the FERARI component RME-1 shows a beads-on-a-string patterns near the apical membrane. Additional signal can be observed at lateral and basal membranes. Most of the GFP-RME-1 signal is lost in knock-downs of FERARI components *(spe-39, vps-45, rabs-5, rfip-2* and *rab-11)*. (E) Knock-down of *snx-1* and its interactor *rme-8* show FERARI-like phenotypes in RAB-11 compartments (similar to FERARI components in Fig. 2D). (F) Quantification of phenotypes shown in (E). (G) Yeast two hybrid interactions of FERARI subunits with SNX-1. SPE-39 shows weak interaction, but this interaction was verified by exchanging the proteins on the plasmids (not shown). Interaction between RABS-5 and SNX-1 is quite strong in this assay. (H) Networks formed by mCherry-SNX-1 (arrow) are lost in *spe-39(RNAi), vps-45(RNAi)* and *rabs-5(RNAi)* intestinal cells.

Our data indicate that FERARI and SNX-1 should both be on sorting endosomes involved in recycling. To corroborate this notion, we analyzed the localization of RME-1 with respect to SNX-1 and found that RME-1 decorates SNX-1 tubules with discrete puncta (Fig. 5B and C, movie S5). This co-localization was dependent on other FERARI members. When we depleted SPE-39 or other FERARI components the association of RME-1 with internal structures was lost, while the plasma membrane pool appeared somewhat less disturbed (Fig. 5D). This effect was not due to RME-1 degradation in the absence of FERARI (Fig. S5). Thus RME-1 depends on binding to the other FERARI components for recruitment to SNX-1 positive sorting endosomes.

Next, we explored the relationship between SNX-1 and FERARI further. SNX-1 interacted specifically with SPE-39 and RABS-5 in Y2H (Fig. 5E, Fig. S1 A, F and H). Likewise, knockdown of SNX-1 or its interacting partner RME-8 ^43, 44^ resulted in enlarged RAB-11 positive compartments, indistinguishable from those formed upon loss of FERARI (Fig. 5E and F compare to Fig. 2D and E). These data suggest that FERARI and SNX-1/RME-8 act in the same pathway (Fig. 1I). More importantly, FERARI(RNAi) led to loss of the tubular SNX-1 structures (Fig. 5H), indicating that FERARI is required for the maintenance of the SNX-1 recycling compartment, which feeds into the RAB-11 recycling pathway.

### The SNAREs SYX-6 and SYX-7 act in the FERARI-dependent RAB-11 recycling pathway

As a multi subunit tethering complex (MSC) FERARI should also have a functional SNARE binding module. The SM protein VPS-45 was shown to interact directly with SNAREs SYX-7 and SYX-16 ^29^. While *syx-7(RNAi)* caused the enlarged RAB-11 compartment phenotype characteristic of FERARI loss, *syx-16(RNAi)* had only a mild effect (Fig. 6A and B). When we explored other SNAREs in the endosomal system, we identified SYX-6 as another potential FERARI SNARE, while VAMP-7 and VTI-1 appeared to have no direct function in RAB-11-dependent recycling (Fig 6A and B). Taken together, our results indicate that FERARI fulfills the formal requirements of a multi subunit complex tether in that it has a SNARE- and a Rab-interacting module, which could be sufficient to bring two membrane entities together to drive membrane fusion.

**Figure 6:**
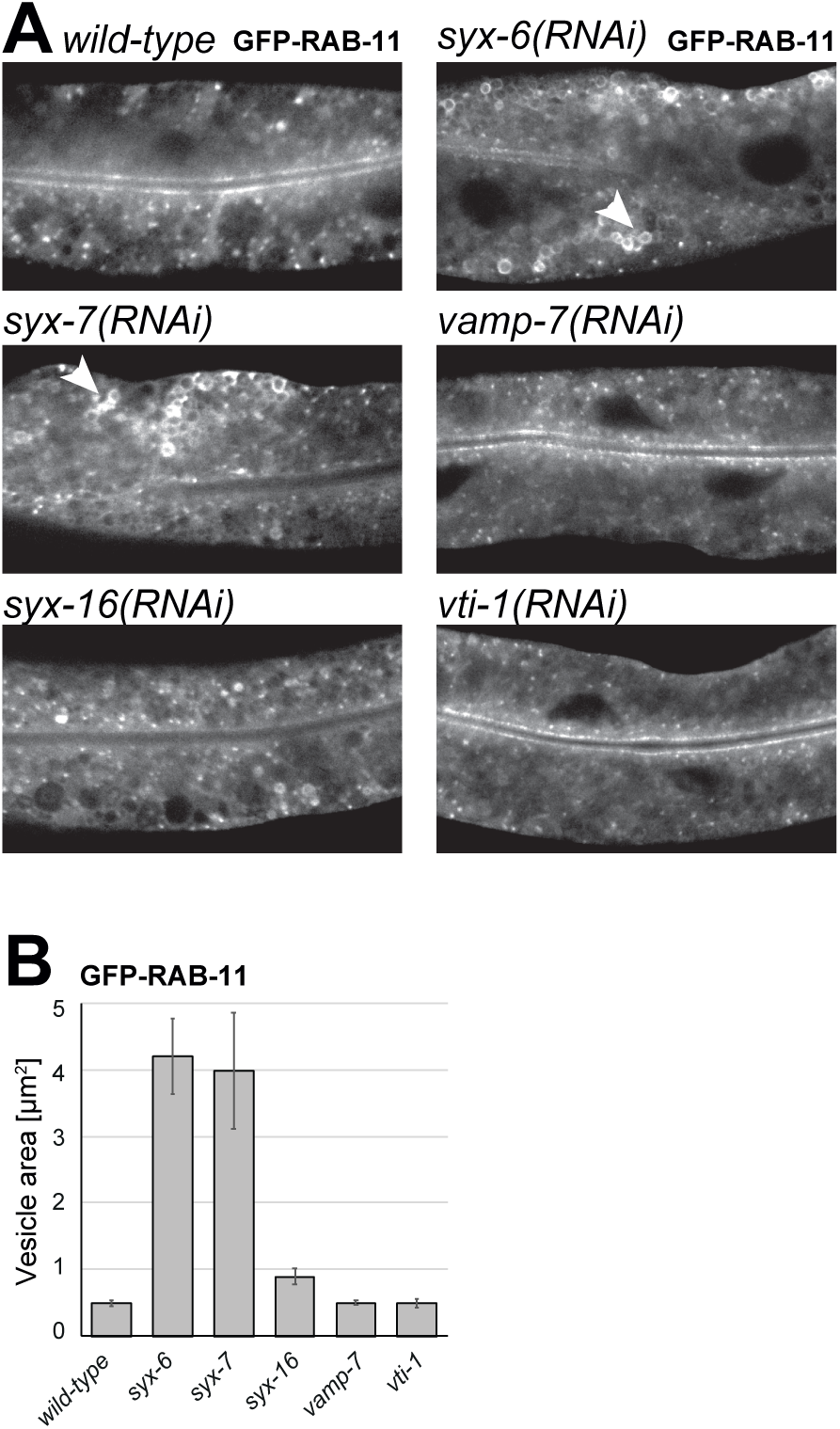
The SNAREs SYX-6 and SYX-7 act in the FERARI-dependent RAB-11 recycling pathway. (A) SNARE knock-downs show distinct phenotypes on RAB-11 compartment. While *syx-6* and *syx-7(RNAi)* show enlarged RAB-11 compartments (arrows) *syx-16, vamp-7* and *vti-1(RNAi)* affect only RAB-10 compartments (not shown) and leave RAB-11 mostly unchanged compared to *wild-type* (lower panels). (B) Quantification of SNARE knock-down phenotypes (6 worms with 10 structures measured for each). Included are SNAREs with RAB-10 phenotype *(syx-16, vamp-7, vit-1)*.

### RAB11-positive endosomes kiss-and-stay at sorting endosomes

But how would a tethering factor at the sorting endosome drive recycling endosome formation? FERARI is special with respect to the different functions it combines. Next to the classical tethering activities, it also contains EHD1/RME-1, which acts as a pinchase^14^. Thus, FERARI may promote membrane fission and fusion at the recycling compartment of sorting endosomes. If this was the case, we may be able to detect the arrival of a RAB-11-positive structure at the recycling compartment of the sorting endosome. We detected RAB-11 positive structures docking onto the SNX-1 positive sorting endosome (Fig. 7A stills of the movies, movies S8-S10). These RAB-11 positive structures are small with low fluorescence intensities (Fig. 7C). The RAB-11 positive structures appear to stay at the SNX-1 positive sorting endosome during the duration of the movie. Yet, we also observed instances in which RAB-11 positive structures were docked on the sorting endosomes and were then released (Fig. 7B, movies S11-S13). Importantly, those RAB-11 endosomes appeared to bigger and brighter (Fig. 7C). We take this as an indication that RAB-11 endosome picked up cargo and lipids at the sorting endosome for subsequent delivery at the plasma membrane, consistent with a kiss-and-stay mechanism.

**Figure 7:**
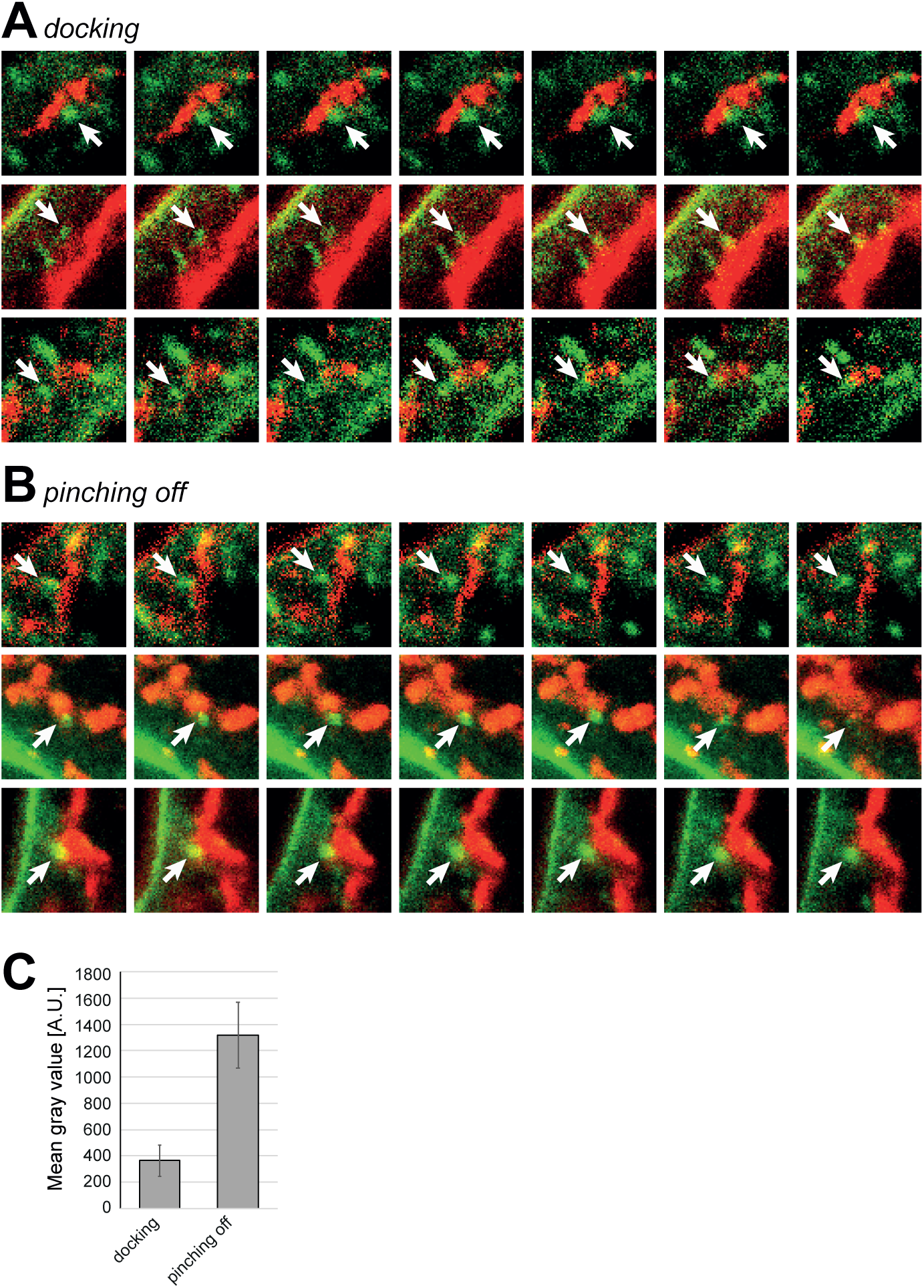
RAB11-positive endosomes kiss and stay at sorting endosomes. (A) Movies with GFP-RAB-11 and mCherry-SNX-1 compartments, showing docking of globular RAB-11 to SNX-1 tubules. Stills from movies are shown, arrows point to RAB-11 compartment of interest. Please note the yellow overlap in later pictures to the right. (B) Stills from movies showing pinching off of RAB-11 globular compartments from SNX-1 tubules. Arrows point to RAB-11 compartment starting out with some overlap (yellow) and moving away from SNX-1 tubules (images to the right). Time between stills is 30 seconds (every 3^rd^ image in movies (see supplementary materials). (C) Docking RAB-11 compartments are less bright than pinching off RAB-11 compartments. Brightness was measured over 10 frames of 5 movies each.

## Discussion

How sorting into recycling endosomes occurs at sorting endosomes still remains enigmatic. Here, we report the existence of a multisubunit tethering complex, FERARI, which plays a role in recycling to the apical and basal lateral domains of polarized cells in *C. elegans* and Rab11-dependent recycling in mammalian cells (Fig. 8A). FERARI represents a tether as it combines a SNARE-interaction module - VPS45-with Rab interaction modules -Rab11FIP-5/RFIP-2 and RBNS5/RABS-5 ^27^. Unlike its relatives, HOPS and CORVET, FERARI tethers membranes of different identities. Another remarkable difference is that FERARI contains a dynamin-like protein EHD1/RME-1. This composition provides FERARI with two rather distinct and opposing activities: membrane fission and membrane fusion.

**Figure 8:**
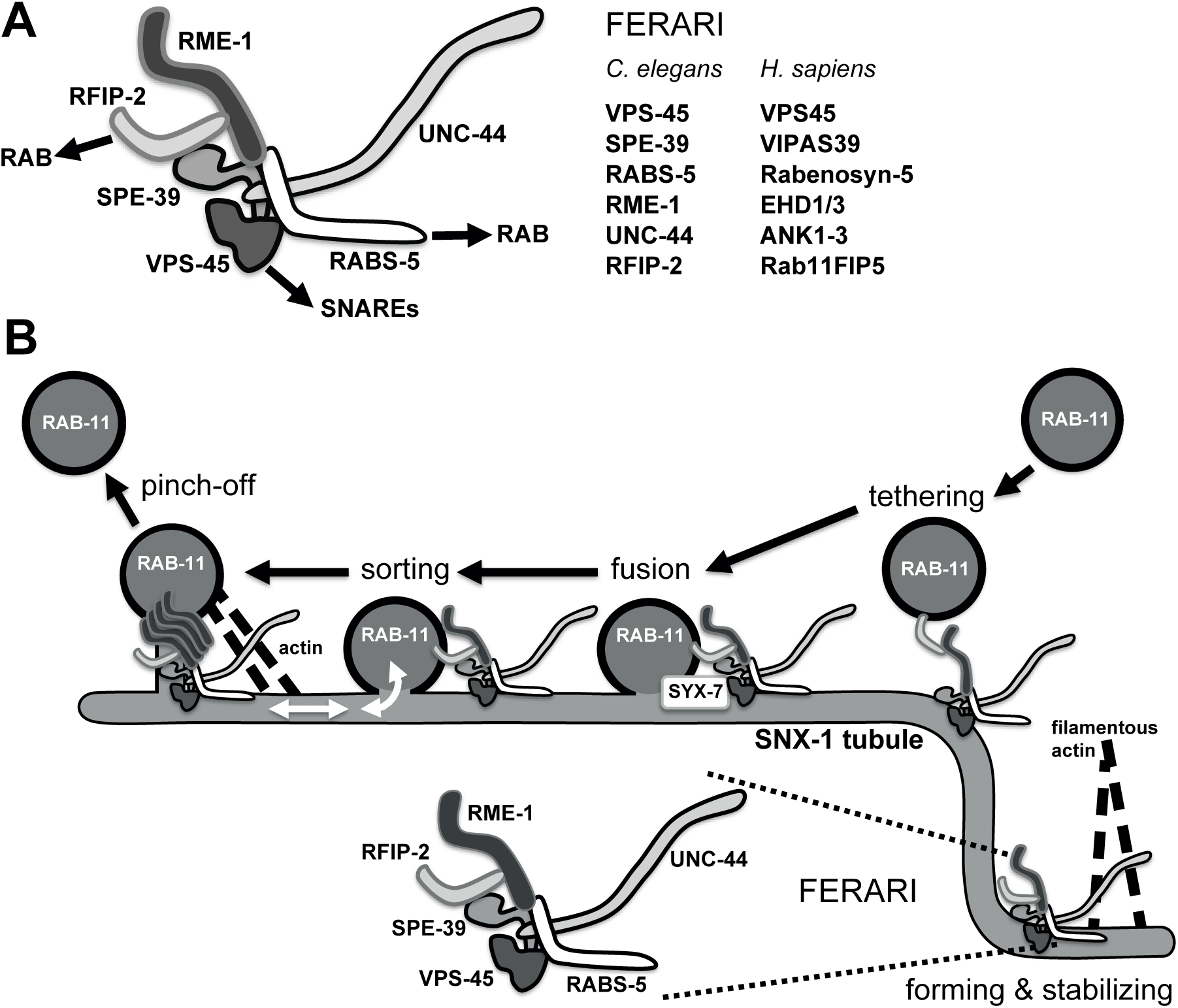
Model of FERARI recycling. A putative structure of FERARI is shown below, based on interactions between the components and approximate size of proteins. Functions of FERARI might include forming and stabilizing SNX-1 sorting/recycling tubules (this might happen near or on RAB-5 positive structures and involve the interaction of UNC-44 with actin). FERARI could tether RAB-11 compartments to these tubules, followed by membrane fusion (mediated by VPS-45 binding of SNAREs SYX-6 and SYX-7). The RAB-11 compartment would be stabilized with an open pore to the SNX-1 tubule because of the presence of an RME-1 constriction ring. After sorting of cargo, the pinching activity of RME-1 (maybe coupled to UNC-44 interaction with actin) will allow the RAB-11 compartment to leave the sorting station.

Based on our data, we propose the following model (Fig. 8B). We assume that FERARI is recruited by RAB5 onto the sorting endosome, where it associates with SNX-1 and helps to stabilize SNX-1 positive structures. At this point we can only speculate about the mechanism. However, ANK1 and ANK3 are known to link membranes to the cytoskeleton and would therefore be well suited to stabilize tubular networks. VPS-45 mediates association of the tethering complex with the SNAREs SYX-6 and SYX-7, while RAB11FIP5/RFIP-2 will engage with RAB11 on a recycling structure to bring it closer to the sorting endosome. FERARI then would induce SNARE complex assembly and promote membrane fusion, similarly to what has been reported for the DSL, HOPS and CORVET complexes ^24, 45, 46^. Even though we anticipate full membrane fusion, our data indicate that the RAB11 positive membrane does not flatten and retains its globular structure. The barrier for the membrane flattening would be provided by EHD1/RME-1. EHD proteins have been shown to assemble in ring structures similar to dynamin ^15, 47^. In this state, cargo to be recycled to the plasma membrane and lipids could diffuse into the RAB11-positive structure. Finally, EHD1/RME-1 would promote scission between the RAB11-positive recycling endosome and the tubular part of the sorting endosomes. Our model corresponds to a kiss-and-stay model and is similar to the kiss-and-run model in neurotransmitter release, except that in our case the docked entity is not releasing but is taking up content. Even the ‘flickering’ in the kiss-and-run model of membrane fusion and fission could be explained with two activities combined in FERARI through fusion promoting function of VPS45 and membrane scission ability of EHD1/RME-1 protein. In neuroendocrine cells, secretory granules maintain their shape and do not flatten out during fusion with the plasma membrane. Instead, they are released again from the plasma membrane in a process referred to as cavicapture or fuse-pinch-linger ^48, 49^. The coordination of fusion and fission events could be mediated by EHD1 through interaction with either VPS45 or another SNARE regulator protein, SNAP29, ^50^. Thus, FERARI provides a platform on sorting endosomes on which fission and fusion of RAB11 positive structures are coordinated. Rabenosyn-5/RABS-5 could potentially act as the scaffold because it can interact with all FERARI members and sorting nexins.

Our model is supported by our finding that the RAB-11 compartments that dock onto the sorting endosome are smaller than the ones that leave. Moreover, most components of the FERARI have been implicated in recycling previously ^17, 21, 26, 28, 51^. Now our data put them into context, providing a temporally and spatially controlled concerted action of the complex. Some of the FERARI components do not show complete co-localization with other FERARI members, suggesting that they might perform additional functions in the cell. Similarly, the HOPS components VPS39 and VPS41 have been reported to have HOPS-independent functions ^52–54^.

EARP is another MTC located on RAB4-positive recycling endosomes ^55^. However, the function of EARP on the RAB4 recycling endosome remains elusive. It is conceivable that these tethers, EARP and FERARI, not only bring membranes together but are also required to build or maintain domains on which recycling endosomes can form. We observed that FERARI not only was involved in RAB11-dependent recycling but also early on in the formation/maintenance of the tubular structures on which sorting would occur. In the case of FERARI, we would postulate that rabenosyn-5 and the ANK proteins might be involved in such domain formation. If this hypothesis is correct, we can expect that the number of tethering complexes on sorting endosomes will increase. In fact, there is at least one candidate for such an additional tether already and that is CHEVI ^27^, which has been proposed to act in recycling as well as in a-granule formation, a process which requires TGN to MVB transport ^28, 56^. So far only the two components, VPS33B and VIPAS39, of this putative MTC have been identified. Yet, given the localization of these proteins, it is likely that CHEVI acts on sorting endosomes. We propose that MTCs could play a role in the formation of domains required for sorting and docking on sorting endosomes but putatively also on other organelles.

## Material and Methods

### Worm husbandry

*C. elegans* was grown and crossed according to standard methods ^57^. RNAi was performed as previously described ^11^. All experiments were carried out at 20°C, and worms were imaged at the young adult stage (with only few eggs). Worm strains and transgenes used in this study: *bIs1[YP170::GFP + rol-6(su1006)]X, pwIs116 [rme-2p::rme-2::GFP::rme-2 3’UTR + unc-119(+)], pwIs429[vha-6::mCherry-rab-7], pwIs72[vha6p::GFP::rab-5 + unc-119(+)], pwIs90[Pvha-6::hTfR-GFP; Cbr-unc-119(+)], pwIs50[lmp-1::GFP + Cb-unc-119(+)], pwIs414[Pvha-6::RFP::rab-10, Cbr-unc-119(+)], pwIs69[vha6p::GFP::rab-11 + unc-119(+)], pwIs782[Pvha-6::mCherry::SNX-1], pwIs87[Pvha-6::GFP::rme-1; Cbr-unc-119(+)], [Pdhs-3::dhs-3::GFP], pwIs481[Pvha-6::mans-GFP, Cbr-unc-119(+)]*.

### Microscopy

Living worms were imaged as described ^11^. Levamisole (50 mM) was used for immobilization and 2% agarose pads were cast onto microscopy slides; worms were sealed under cover slips using Vaseline. Images were acquired using a spinning disk confocal system Andor Revolution (Andor Technologies, Belfast, Northern Ireland) mounted onto an IX-81 inverted microscope (Olympus, Center Valley, PA), equipped with iXon^EM^+ EMCCD camera (Andor Technologies). A 63× 1.42 N.A. oil objective was used, where each pixel represents 0.107 μm. Solid-state 488 nm and 560 nm lasers were used for excitation and exposure time was 100 ms. Images were averaged 4 times for oocytes and 32 times for intestine and coelomocytes. Alternatively, an Olympus Fluoview FV3000 system with a high sensitivity-spectral detector (HSD) was used (PTM voltage= 500). The objective was 60x with silicone oil, and the Galvano scan device was used. Pixel size corresponds to 0.098 μm. Sampling speed was 8.0 μs/pixel, zoom= 2.1, lasers 488 (GFP) or 561 (RFP, mCherry) were set at 4-10%. All images were processed in the same way for corresponding experiments.

Images of SNX-1 tubules with different GFP markers were obtained on a Zeiss LSM 880 microscope with Airyscan. The fast mode was used and images were processed using the Zen Black software. Worms were treated with 20 mM sodium azide to avoid any movement during image acquisition.

### Compartment quantifications

The length of RAB-10 tubules was measured by applying a freehand line ROI in one z plane, for size of RAB-11 compartments the elliptical selection tool was used and the area in pixels was measured. Both methods might lead to an underestimate of the real sizes, because tubules and globular compartments were not measured in 3D projections, but the differences between wild-type and mutant phenotypes were sufficiently large so that more complicated quantification methods were not essential. In all cases, 6 different worms were analysed and 10 structures per worm were measured (in different z planes).

The size of RAB-10 globular compartments could be measured in a more automated way because they are very bright and sufficiently separated from each other (in contrast to RAB-11 where only the outlines of the enlarged compartments were visible and vesicles clustered together). Maximal z projections were carried out on the whole stack of pictures, yielding about 50-120 vesicles per worm (6 worms per condition were measured). Threshold was set and the pictures were transformed into binary images. The “close-“ and “watershed” function was applied to get rid of single pixels and fused objects. Particles were analysed (size setting= 0-infinity and circularity= 0.5-1.0). Extremely large accumulations of objects that escaped the “watershed” function were manually excluded from the final analyses.

### Interaction studies with worm proteins

*C. elegans* proteins SPE-39 and VPS-45 were expressed in *E. coli* Rosetta (DE3) cells. GST-tagged SPE-39 was obtained by cloning the worm cDNA into pETGEXCT vector, while His-tagged VPS-45 cDNA was cloned into pRSF-2 Ek/LIC vector. 500 ml *E. coli* cultures bearing both plasmids (grown in LB+amp+kan) were induced with 0.5 mM IPTG overnight at 37°C and collected in buffer with 50 mM HEPES pH 7.7, 300 mM NaCl, 10% glycerol and 1% TritonX 100 (with protease inhibitors) for purification. Cells were lysed with 1 mg/ml lysozyme, 20 μg/ml DNaseI and 2x 30 sec sonication. After a 16,000 g spin, the cleared lysate was incubated overnight with Ni-NTA beads. Beads were washed 4x and eluted with 250 mM Imidazole. The eluate was pooled and bound overnight to glutathione beads. Worm Lysate from 100 ml liquid culture (with many adults) was prepared in the same buffer (with glass beads in a FastPrep machine for 2x 30 sec). The worm extract was centrifuged at 16,000 g for 30 min to yield a clear worm lysate that was added to the washed Glutathione beads with bound SPE-39/VPS-45 complex. Proteins were allowed to bind overnight and finally the beads were washed 4x with lysis buffer and 4x with lysis buffer lacking any kind of detergents for mass spectroscopy analysis. Proteins were kept at 4°C at all times.

Yeast two hybrid assays were carried out using worm cDNA cloned into the vectors pEG202 (with DNA binding domain DBD) and pJG4-5 (with activation domain AD). Fastest results were obtained by Gibson Assembly cloning (NEBuilder, NEB; E55205). pEG202 clones also obtained a 3xFLAG tag at the C-terminus for pull-down experiments. Growth assays were carried out with 6 colonies from yeast transformation (strain EGY48+pSH18-34 plasmid for LacZ expression) for 3-7 days at 30°C on plates lacking Leucine and with addition of 2% galactose for induction of pJG4-5 expression. For pull-down experiments, 50 ml cultures of yeast were induced with 2% galactose at OD600= 1.0 for 6 hours. Cells were lysed in 50 mM HEPES pH 7.7, 300 mM NaCl, 10% glycerol and 1% TritonX 100 (with protease inhibitors) in a FastPrep device (2x 30 sec). 25 μl of magnetic beads (slurry) were added for binding of tagged proteins (either anti-HA (Pierce; 88837) or anti-FLAG beads (Sigma; M8823)) and rotated overnight. After 4x washing, proteins were eluted with Laemmli sample buffer at 95°C for 5 min and applied to SDS-PAGE gels for immunoblot analysis with FLAG and HA antibodies (FLAG: Sigma; F3165, HA: Sigma HA-7 clone; H9658).

### Cell culture, transfection and CRISPR/CAS9 knockout in mammalian cells

HEK293 and HELA cells were cultured and maintained in DMEM (Sigma) high glucose medium with 10% FCS (Bioconcept), penicillin–streptomycin (1%), Sodium pyruvate and L-glutamine. Cells were plated 1 d before transfection at 60-70% confluency and later transfected for 48 hours using Turbofect transfection reagent (Thermo Scientific; R0532) according to the manufacturer’s instructions. 2-5 μg of DNA was used per reaction based on a 10 cm dish.

For CRISPR/Cas9-mediated knockout, guide RNAs were selected using the CRISPR design tool (http://chopchop.cbu.uib.no/). Two guide RNAs were designed from two different exons for each target gene. Annealed oligonucleotides were cloned into two different plasmids containing GFP marker and Puromycin resistance, respectively. In brief, HELA cells were seeded at 2 × 10^6^ cells per 10 cm dish. The following day, cells were transfected with 2.5 μg of the plasmids (control vectors without insert or vectors containing a guide RNA against target gene). Transfecting media was exchanged with fresh media after 4 h. Cells were treated with puromycin for 24 h after transfection followed by FACS sorting (for GFP+ cells) on next day. For FACS sorting after 48 h of transfection, cells were trypsinized and resuspended in cell-sorting medium (2% FCS and 2.5 mM EDTA in PBS) and sorted on BD FACS AriaIII Cell Sorter. GFP-positive cells were collected and seeded in a new well.

### Immunoprecipitation assays

HEK-293 cells were co-transfected with mentioned DNA constructs. After 36-48 h of transfection, protein extracts were prepared in lysis buffer (1% NP-40, 50 mM Tris/HCl pH 7.5, 150 mM NaCl) and Halt protease inhibitor cocktail (Thermo Scientific; 186 1279) at 4°C for 20 min followed by centrifugation at 4°C for 20 min at 13,000 rpm. Immunoprecipitations were performed as previously described ^58^. In brief, protein extracts were incubated with trap beads (nano bodies for GFP (GFP-Trap^®^_A; gta-20-chromotek), myc (Myc-Trap^®^_A; yta-20-chromotek), turbo-GFP (TurboGFP-Trap_A; tbta-20-chromotek)) for 4 h at 4 °C with rotation, and then washed 5x with lysis buffer (1 ml). Protein complexes were eluted by heating beads for 5 min at 95°C in 2x sample buffer and resolved by SDS-PAGE on 10 and 12.5% gels followed by immunoblot analysis. Blots were developed using Amersham ECL Prime Western Blotting Detection Reagent (RPN2236) and X-ray film (Amersham Hyperfilm ECL-28906839).

### Western blot analysis

Cells were collected and lysed in lysis buffer (50mM Tris/HCl, 150 mM NaCl, 1% NP-40) containing a protease inhibitor cocktail (Roche). Protein concentrations were determined in all experiments using the Bio-RAD protein assay (Bio-RAD, 500-0006) and 20-40 μg of total protein was loaded onto either 10 or 12.5 % SDS-PAGE before transfer onto nitrocellulose membranes (Amersham Protran; 10600003). Membranes were blocked 5% milk, 0.1% Tween20 for 60 min at RT. First antibody incubation was O/N at 4°C and the secondary HRP coupled antibodies incubated for 1 h at RT. The blots were developed using Amersham ECL Prime Western Blotting Detection Reagent (RPN2236) and X-ray film (Amersham Hyperfilm ECL-28906839) or the Fusion FX7 (Vilber Lourmat) image acquisition system.

### Antibodies

The antibodies used in this study: polyclonal rabbit anti-VIPAS39 (20771-1-AP; proteintech; 1:2000), polyclonal rabbit anti-Rab11-FIP5 (NBP1-81855; Novus Biologicals;1:2,000), polyclonal rabbit anti-EHD1 (NBP2-56035; Novus Biologicals; 1:2,000), polyclonal rabbit anti-rabenosyn-5 (NB300-813; Novus Biologicals; 1:2,000), monoclonal mouse anti-myc (9E10) (1:3,000 for WB and 1:200 for Immunostaining; Sigma-Aldrich; M4439), polyclonal rabbit anti-GFP (TP401; Torrey Pines; 1:3,000 for WB and 1:200 for Immunostaining). For pulldowns Trap beads (Nano bodies) were used. GFP-Trap_A (chromotek, gta-20) for GFP pulldowns and myc-Trap_A (chromotek; yta-20) for myc pulldowns. HRP-conjugated goat anti-Mouse IgG (H+L) Secondary Antibody (Thermo Fisher scientific; 31430; 1:10,000), polyclonal HRP conjugated goat-anti-rabbit IgG (Thermo Fisher scientific; 31460; 1:10,000) were used (incubated for 1 h at RT) to detect bound antibodies with and an ECL system (ECL prime, Amersham, RPN2232). Alexa Fluor 488-goat anti-rabbit IgG (H+L) (Invitrogen; A-11034) and Alexa Fluor 594-goat anti-mouse IgG (H+L) Cross-Adsorbed Secondary Antibodies (Invitrogen; R37121) were used for immunofluorescence.

### Immunostaining in mammalian cells

Cells were plated onto sterile 13 mm glass coverslips. Cells were fixed with 2% paraformaldehyde for 15 min, permeabilized (0.1% TritonX 100 in PBS) for 5 min and blocked with 2% BSA containing 5% goat serum in PBS for 1 h. Cover slips were incubated in Primary antibodies for 2 h and washed in PBS for 5x followed by 1 h incubation in fluorescently-tagged secondary antibodies. After secondary antibody incubation, cover slips were washed again 5x in PBS and mounted onto glass slides using Fluoromount-G (Southernbiotech;0100-01). Images were taken with an inverted Olympus FV1000 confocal microscope using a Plan Apochromat N 60x/1.40 silicon oil objective with z-stacks. Co-localization studies were performed using the ImageJ co-localization plugin JACoP.

### Live cell imaging

For live imaging, cells were plated in 8 well chambered coverglass and media was replaced with warm imaging buffer (5 mM dextrose (D(+)-glucose, H2O, 1 mM CaCl2, 2.7 mM KCl, 0.5 mM MgCl2 in PBS) just before imaging. Images were taken at 37°C on an inverted Axio Observer Zeiss microscope (Zeiss, Oberkochen Germany) using a Plan Apochromat N 63×/1.40 oil DIC M27 objective with a Photometrics Prime 95B camera. Z-stack images were processed by using the OMERO client server web tool and Fiji.

### Quantification of Rab11-positive endosomes

Segmentation and analysis were performed on manually chosen regions of interest (ROI) using a custom script for Fiji^59^ as follows: 1) A 3D White Top-Hat filter ^60^ was applied to the original image in order to homogenize the background and used to compute 3D seeds^61^ with subpixel accuracy. 2) Objects were segmented on the original image using an iterative threshold^62^ and converted to labels. 3) Touching objects were then separated by a 3D watershed^63^ using the previously identified seeds on the label image. 4) The resulting image was then added to the 3D ROI Manager^63^ to exclude remaining laterally touching objects and finally perform intensity and size measurements per object. 1,500-2,500 Rab11-positive particles were analyzed from 45-60 cells for each condition. The script is available upon request.

### DNA plasmid sourcing

Commercially available Plasmids were obtained from Addgene and Origene. VPS33B-Myc-DDK (Origene; RC203870), Myc-DDK-VIPAS39 (Origene; RC202264), GFP-VIPAS39 (Sino.Bio; HG22032_ACG), Myc-DDK-VPS45 (Origene; RC206027), Turbo-GFP-VPS45 (Origene; RG206027), Myc-DDK-rabenosyn-5 (Origene; RC239555), GFP-rabenosyn-5 (Addgene; 37538), RFP-rabenosyn-5 (Addgene; 37537), GFP-RAB11 (Addgene; 12674), Myc-DDK-ANK1 (Origene-RC220763), Myc-DDK-FIP5 (Origene; RG206173), Myc-DDK-EHD1 (Origene; RC211158), Myc-DDK-FIP2 (Origene; RC210414). Plasmids Px458 GFP (Addgene; 48138) and Px459 Puro (Addgene; 62988) were used for cloning gRNAs.

### RNA isolation, cDNA preparation and quantitative real-time PCR (qPCR)

Total RNA was collected from human cells by using RNeasy mini kit (QIAGEN_74104) according to the manufacturer’s protocol. Afterwards, 1000 ng RNA was reverse transcribed into cDNA using cDNA Synthesis Kit GoScript™ Reverse Transcriptase and GoScript™ buffer mix oligo-dT (Promega_A2791) following the instructions given by the suppliers. 250 nM primers and three microliter of the cDNA product was then used for Quantitative real-time-PCR using GoTaq^®^ qPCR Master Mix (Promega_A600A) supplemented with CXR Reference Dye (Promega_C541A) that was then analyzed using an StepOnePlus™ Real-Time PCR System PCR machine (Applied Biosystem). The relative levels of the target transcripts were determined using the human beta actin transcript as reference.

## Acknowledgements

We wish to thank Sheuli Begum and Jonas Fürst for excellent technical assistance with some of the experiments. The proteomics analysis was performed in the Proteomics Core Facility of the Biozentrum with the help of Suzette Moes and Paul Jenö. The Imaging Core Facility of the Biozentrum facilitated the generation of movies and in particular Laurent Guerard provided support for image analysis. Cells were sorted in the FACS Core Facility of the Biozentrum. We are grateful to Ian G. Macara for critical comments on the manuscript. Barth Grant and Pingsheng Liu are acknowledged for sharing strains. Some strains were provided by the CGC, which is funded by the NIH Office for Research Infrastructure Programs (P40 OG010440). This work was supported by the Swiss National Science Foundation (CRSII3_141956, 31003A_141207) and the University of Basel.

---

**Suppl. Figure 1:**
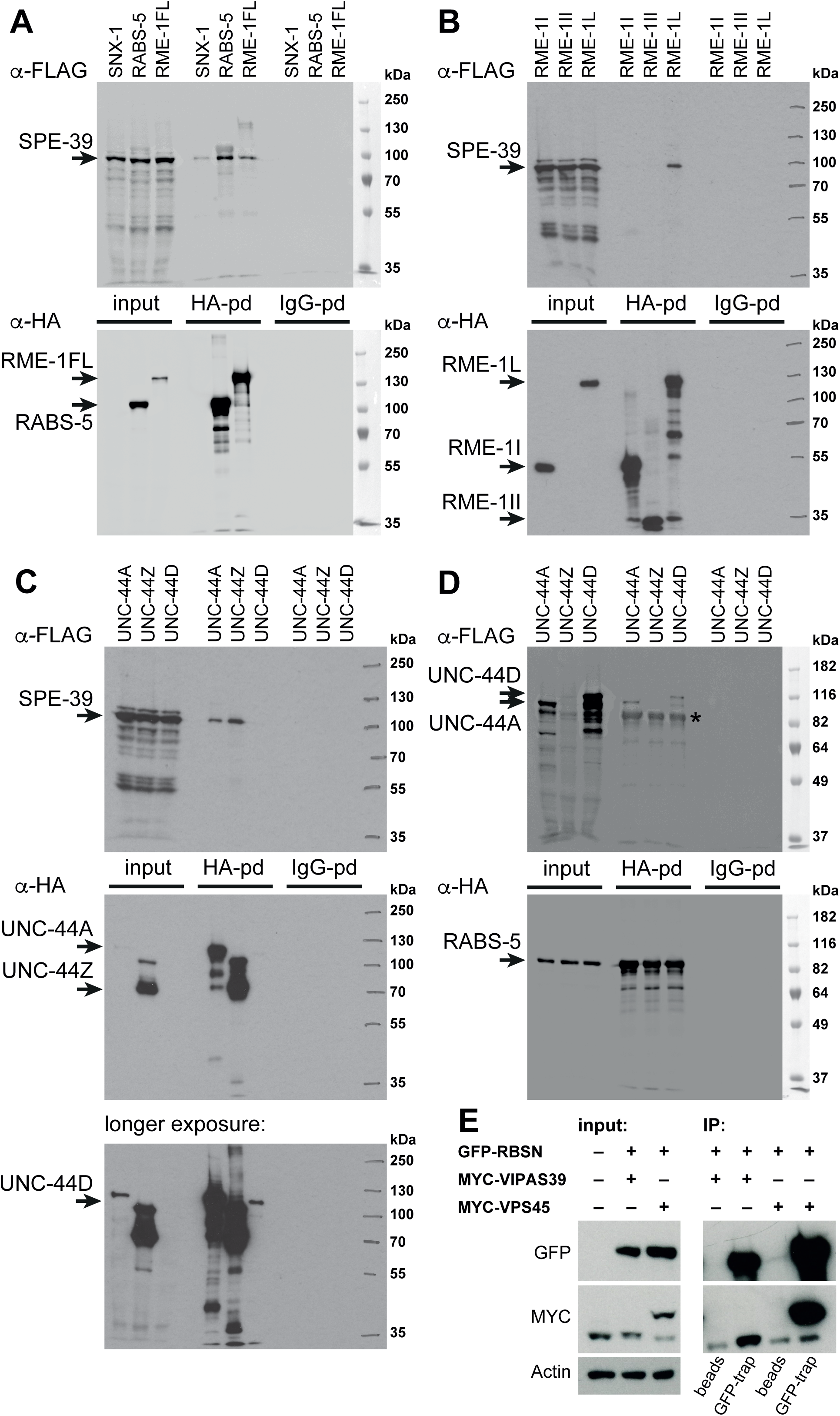
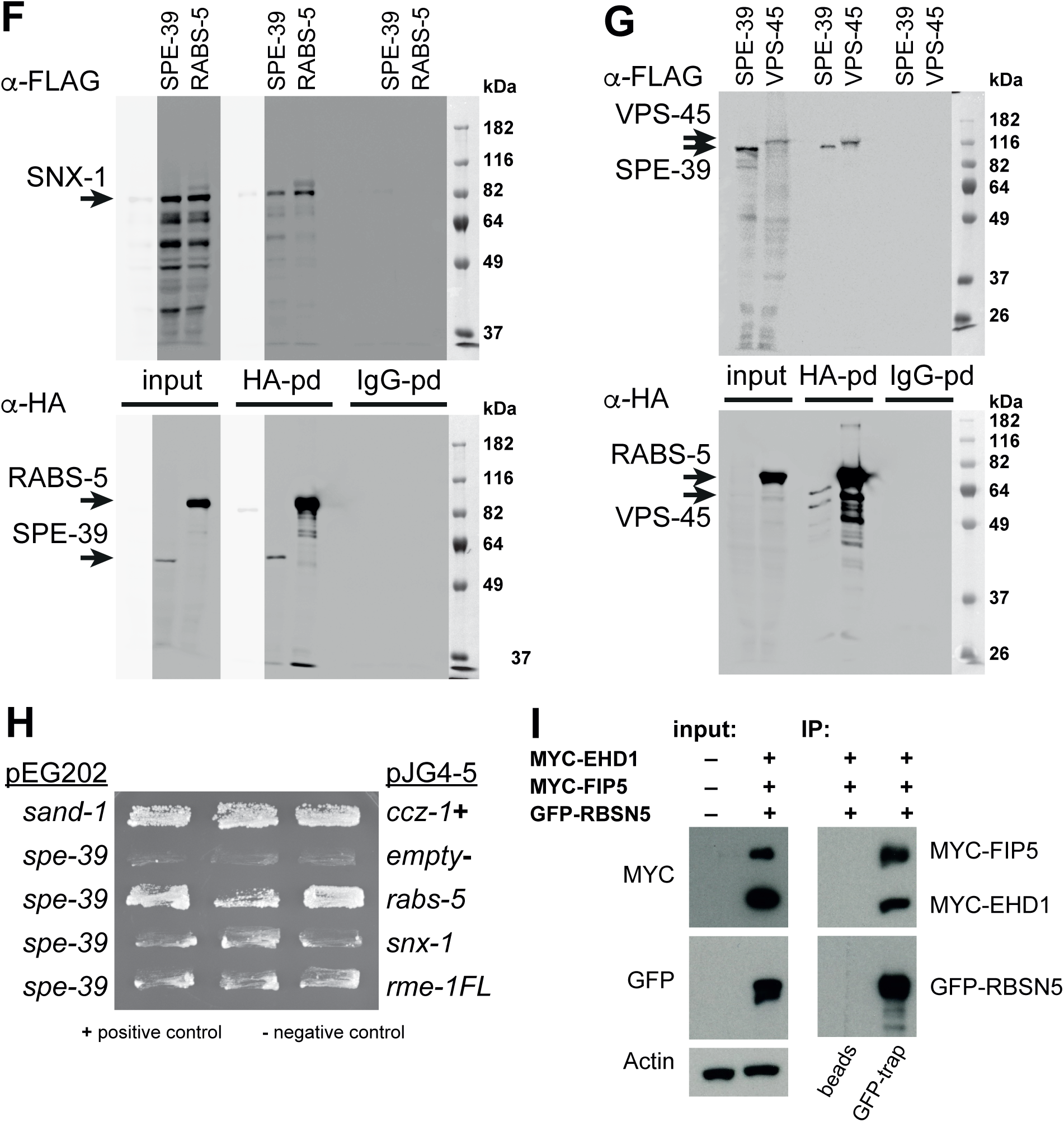
Pull-downs of FERARI subunits show multiple interactions in the complex. (A) Pull-downs were performed with extracts form yeasts expressing SPE-39-FLAG and the proteins indicated at the top (with HA tag). Control pull-downs with beads coupled to IgG were also carried out. Arrows on the left indicate the sizes of expressed proteins. SNX-1 was not expressed in these experiments (but it could be analyzed when cloned into the pEG202 plasmid). (B) Pull-downs showing interactions between SPE-39-FLAG and HA-RME-1 fragments. (C) Pull-downs experiments using SPE-39-FLAG and fragments of UNC-44 (with HA tag). UNC-44D fragment expression is low and can be seen on longer exposure as indicated. (D) Pull-downs showing interaction between HA-RABS-5 and fragments of UNC-44 (coupled to FLAG). UNC-44Z was not expressed, anti-FLAG antibody cross-reacted with RABS-5 (grey bands indicated by asterisk). (E) Western blot of the interactions between transiently over-expressed human FERARI components rabenosyn-5, VIPAS39 and VPS45. GFP-Rabenosyn-5 was precipitated with GFP-trap beads, and myc-VIPAS39 and myc-VPS45 co-precipitated. n= 3 independent experiment (F) Pull down experiment showing the interaction of SNX-1 with SPE-39 and RABS-5. (G) Interactions between SPE-39 and VPS-45 as well as VPS-45 and RABS-5 in the yeast two hybrid system expressing worm proteins. (H) Yeast two hybrid growth assay showing interactions between SPE-39 bait and RABS-5, SNX-1 and RME-1 prey proteins). (I) Western blot depicting the interactions between human protein Rabenosyn-5 with EHD1 and Rab11FIP5, co-expressed in HEK-293 cells. GFP-Rabenosyn-5 was used as bait to co-immunoprecipitate myc-tagged EHD1 and Rab11FIP5. n= 3 independent experiments.

**Suppl. Figure 2:**
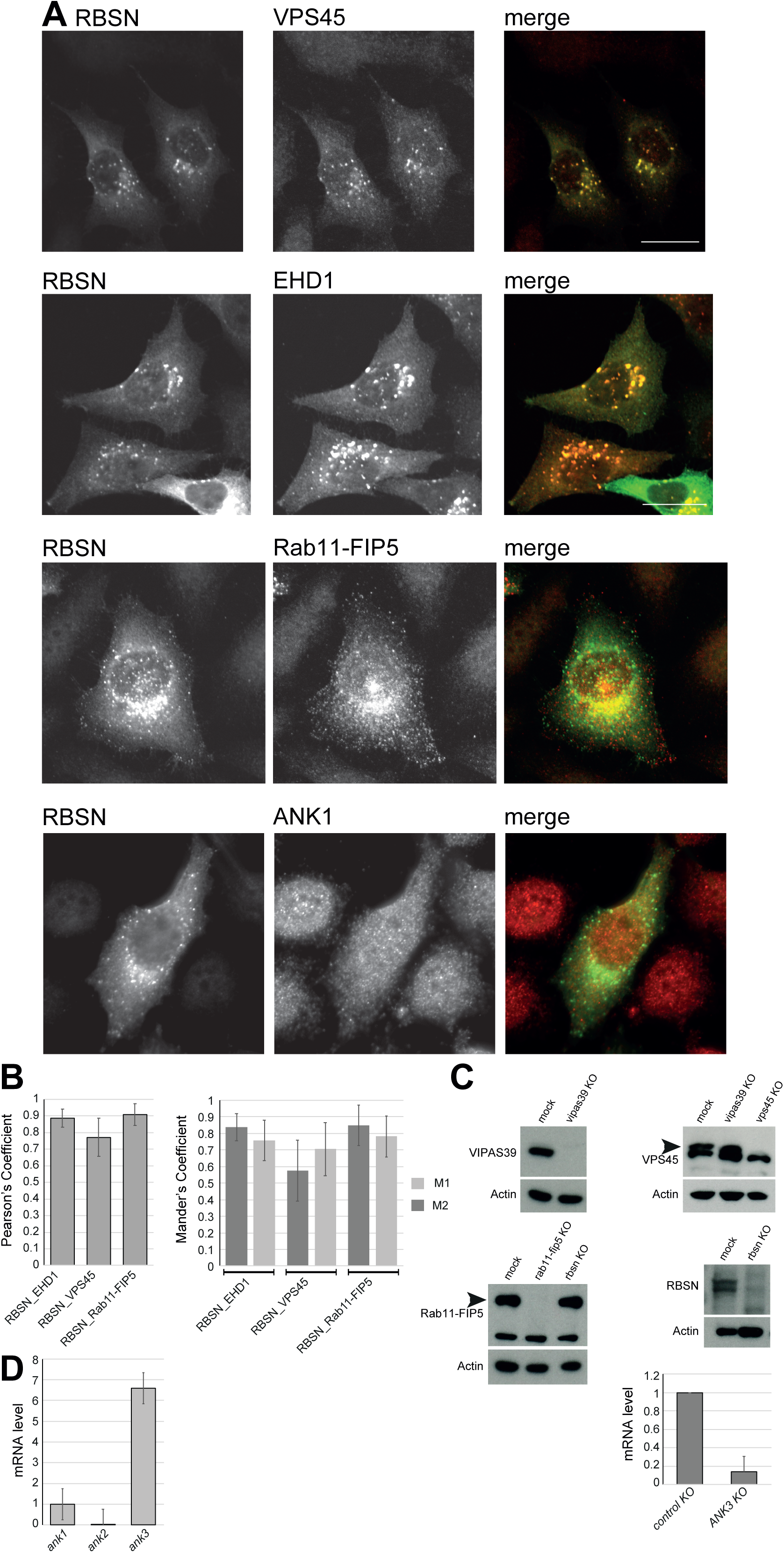
Human FERARI components interact and co-localize in HeLa cells. (A) Co-localization of rabenosyn-5 with other human FERARI components VPS45, EHD1, Rab11FIP5 and ANK1. HeLa cells co-transfected with GFP-Rabenosyn 5 and myc-VPS45, myc-EHD1, myc-Rab11FIP5, myc-ANK1 were subjected to immunofluorescence staining with GFP and myc antibodies. (B) Quantification of co-localization shown in (A). n=30-40 images from 4 independent biological replicates. (C) Western blot depicting the efficiencies of CRISPR-Cas9 knock-out of indicated proteins in HeLa cells used for analyses in Fig. 2A and 3C. The low mRNA level of ank3 indicates that ank3 was knocked out efficiently (D) RT-qPCR data represents levels of ank1, ank2 and ank3 mRNA in HeLa cells at endogenous level. n= 3 independent experiments.

**Suppl. Figure 3:**
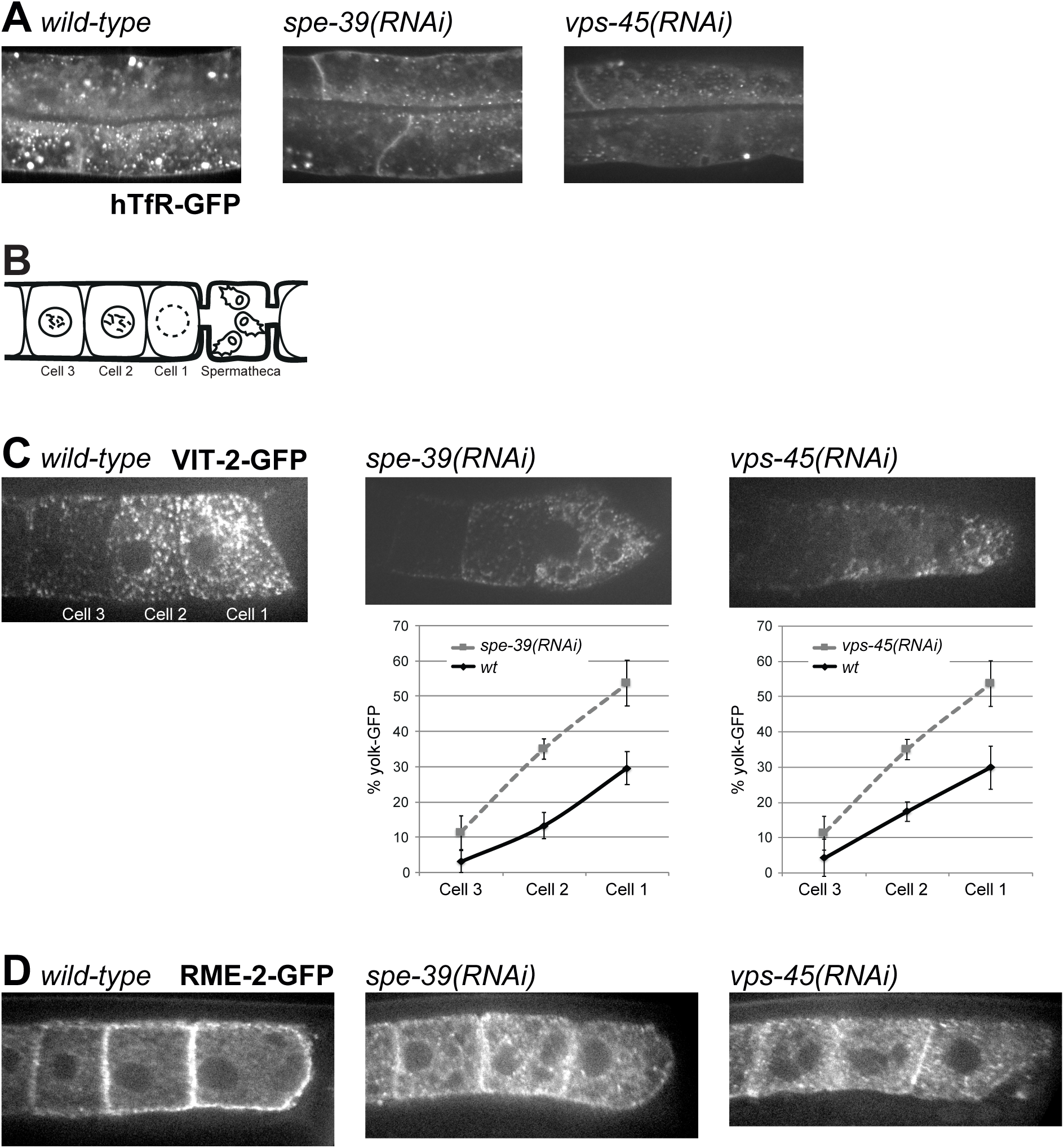
Function of *spe-39* and *vps-45* in endosomal recycling. (A) Recycling of hTfR-GFP is abolished in *spe-39(RNAi)* and *vps-45(RNAi)* worms. Recycling of hTfR-GFP in *wild-type* worms can be seen mainly in the presence of small compartments near the apical membrane (left). Since this receptor has no function in worms, it is also degraded and can be found in larger, more basally located compartments that coincide with lysosomes. Most hTfR-GFP signal is lost in *spe-39(RNAi)* and *vps-45(RNAi)* worms (middle, right). This indicates a loss of recycling activity. (B) Schematic representation of the last 3 oocytes prior to the spermatheca in the worm gonad. (C) Yolk uptake defect of *spe-39(RNAi)* and *vps-45(RNAi)* worms. Wild-type uptake of VIT-2-GFP was quantified in the first 2 oocytes before the spermathecal (schematic in B). The other measurements were normalized to the total fluorescent signal of all 3 *wild-type* cells. Knock-down of *spe-39* and *vps-45* show a reduction of yolk uptake. (D) The yolk receptor RME-2-GFP is mis-localized in *spe-39(RNAi)* and *vps-45(RNAi)* worms (middle, right). In *wild-type* RME-2-GFP signal is found mainly at the plasma membrane, where it binds VIT-2 and is taken up into the cell via endocytosis (left). Knock-down of *spe-39* and *vps-45* causes the loss of RME-2-GFP at the cell surface and concomitant appearance of internal compartments. This observation explains the yolk uptake defect seen above (C).

**Suppl. Figure 4:**
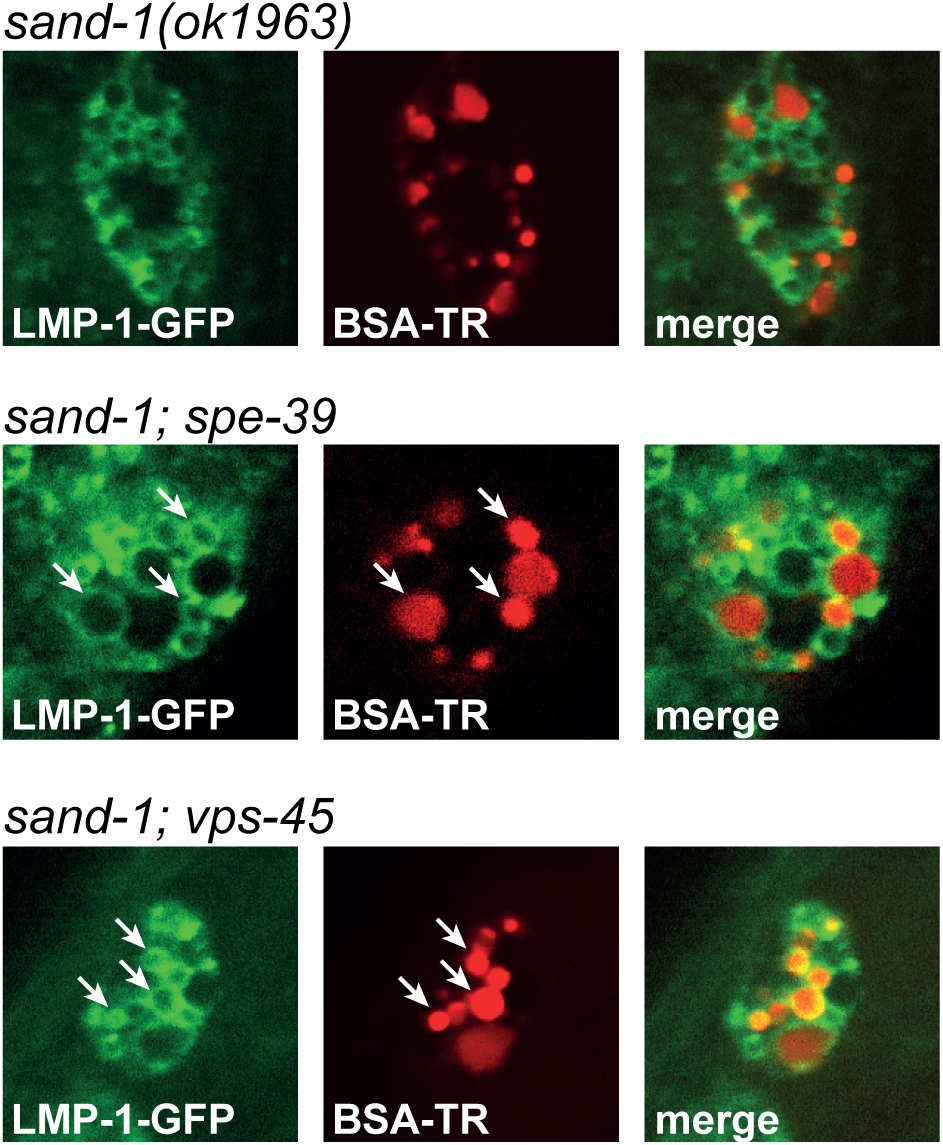
Loss of recycling by *spe-39(RNAi)* or *vps-45(RNAi)* causes more traffic in the degradative pathway. Knock-down of spe-39 or vps-45 induces the degradation pathway in coelomocytes. In *sand-1(ok1963)* coelomocytes BSA-TR is not transported to the lysosomes (LMP-1 compartment) (top images). RNAi of *spe-39* or *vps-45* will favor the transport of BSA-TR to coelomocytes, probably by abolishing recycling (middle and lower images). Arrows indicate LMP-1-positive compartments containing BSA-TR.

**Suppl. Figure 5:**
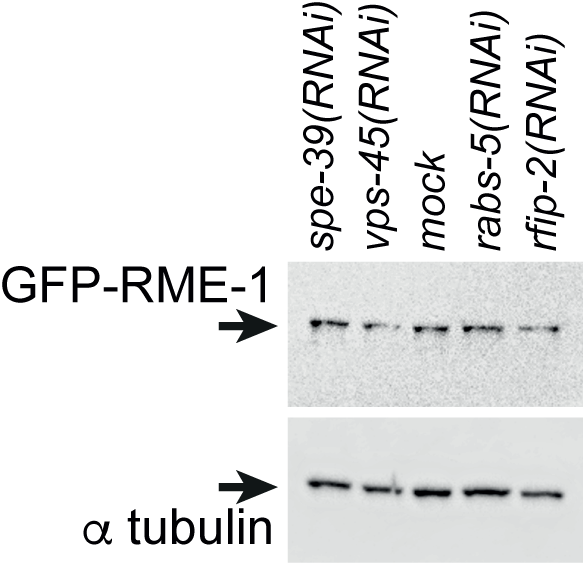
Expression of RME-1 is unaffected by FERARI knock-downs. Protein levels of worms treated with the indicated RNAi. GFP-RME-1 levels were unchanged indicating that the loss of signal in Fig. 5D is due to a recruiting defect and not a loss of protein.

